# Nuclear retention of pre-mRNA involving Cajal bodies during meiotic prophase in plants

**DOI:** 10.1101/2021.04.19.440419

**Authors:** Magda Rudzka, Malwina Hyjek-Składanowska, Patrycja Wróblewska-Ankiewicz, Karolina Majewska, Marcin Gołębiewski, Marcin Sikora, Dariusz Jan Smoliński, Agnieszka Kołowerzo-Lubnau

**Affiliations:** Department of Cellular and Molecular Biology, Nicolaus Copernicus University, Lwowska 1, 87-100 Torun, Poland; Centre For Modern Interdisciplinary Technologies, Nicolaus Copernicus University, Wilenska 4, 87-100 Torun, Poland; Department of Plant Physiology and Biotechnology, Nicolaus Copernicus University, Lwowska 1, 87-100 Torun, Poland

**Keywords:** mRNA nuclear retention, mRNP, Cajal bodies, retained introns, *Larix decidua*, Sm proteins, meiotic prophase

## Abstract

Gene regulation ensures that the appropriate genes are expressed at the proper times. Nuclear retention of incompletely spliced or mature mRNAs emerges as a novel, previously underappreciated layer of post-transcriptional gene regulation. Studies on this phenomenon indicated that it exerted significant impact on the regulation of gene expression by regulating export and translation delay, which allows synthesis of specific proteins in response to a stimulus, e.g. under stress conditions or at strictly controlled time points, e.g. during cell differentiation or development. Here, we found that transcription in microsporocytes, during prophase of the first meiotic division, occurs in pulsatile manner. After each pulse, the transcriptional activity is silenced, but the transcripts synthesized at this time are not exported immediately to the cytoplasm, but are retained in the nucleoplasm and Cajal bodies (CBs). In contrast to nucleoplasm, mature transcripts were not found in CBs. Only non-fully-spliced transcripts with retained introns were stored in the CBs. Retained introns are spliced at precisely defined times, and fully mature mRNAs are released into the cytoplasm, where the proteins are produced. These proteins are necessary for further cell development during meiotic prophase. Our findings provide new insight into the regulatory mechanisms of gene expression based on mRNA retention in the nucleus during the development of generative cells in plants. Similar processes were observed during spermatogenesis in animals. This indicates the existence of an evolutionarily conserved mechanism of gene expression regulation during generative cells development in Eukaryota.

## Introduction

The life course of mRNA begins with transcription by RNA Pol II, splicing, and processing 5’ and 3’ ends, which generally occur at the nuclear sites of transcription, and ends with cytoplasmic translation and degradation. mRNAs are thought to predominantly reside in the cytoplasm for the majority of their lifetime (Halpern. et al. 2015). However, in many cell types, it has been observed that a significant portion of polyadenylated transcripts (approx. 30%) are nuclear retained and undetected in the cytoplasm (Herman et al. 1976, Wagner and Müller-McNicoll 2018). It is now known that some of these transcripts represent the so-called long non-coding RNAs-mRNA-like - that, like the true mRNAs are spliced and polyadenylated, and perform various types of regulatory functions (Inagaki et al. 2005). In recent years, there were reports concerning both animals and plants, revealing that the protein-coding transcripts might be retained in cell nucleus for most of their lifetime (e.g. Halpern et al. 2015, Averbeck et al. 2005, Boothby et al. 2013). Since mRNAs are transcribed and processed at the sites of transcription and translated in the cytoplasm, this lengthy retention period points at possible overlooked role of cell nucleus (Halpern et al.2015). Studies on nuclear retention of mRNAs indicated that it had a significant impact on the regulation of gene expression by regulating the export and translation delay, which allowed the synthesis of specific proteins in response to a stimulus (eg. under stress conditions) or at strictly controlled timepoints. Regulation of gene expression by nuclear retention of mRNAs had been demonstrated, among others, in mammalian metabolic tissues (Halpern et al. 2015), in generative cells (Averbeck et al. 2005, Boothby et al., 2013) or in cells under stress conditions (Prasanth et al. 2005, Ninomiya et al. 2011), and although the final result of such regulation is always the same (i.e. delayed translation), the transcripts can be retained in different forms. In generative cells a regulatory mechanism of gene expression, based on the retention of not fully mature mRNA was observed. Recent studies on male cells of fern *Marsilea vestita* showed that retention and subsequent removal of introns from pre-mRNA regulates the translation pattern during post-transcriptionally regulated spermatogenesis (Boothby et al. 2013). As sequencing analyzes revealed, only one intron was usually left during the mRNA splicing, its length generally not exceeding 100 nt. This intron was removed post-transcriptionally, when a cell was at developmental stage when the protein encoded by the mRNA was needed. This process is a functional mechanism that prevents premature translation of proteins required during specific stages of the development of transcriptionally silent gametes.

In metabolic tissues of mammals (beta cells, liver, and gut) a wide range of protein-coding genes was identified for which the levels of spliced polyadenylated mature mRNAs were higher in the nucleus than in the cytoplasm (Halpern et al. 2015). These included genes such as the ChREBP transcription factor, Nlrp6, glucokinase, and glucagon receptor. A single-molecule *in situ* technique used to quantify nuclear mRNA lifetimes revealed that transcripts of these genes could spend hours in the nucleus before being exported to the cytoplasm. Mammalian genes are transcribed in bursts leading to temporal fluctuations in cellular mRNA levels and variability among identical cells. Nuclear retention can effectively buffer these fluctuations, facilitating lower variability in cytoplasmic mRNA levels. A mechanism of mRNA nuclear retention by "closing" protein coding sequence *mCAT2* (***<u>m</u>**ouse **<u>c</u>**ationic **<u>a</u>**mino acid **<u>t</u>**ransporter **<u>2</u>***) in a long non-coding RNA – CTN-RNA (Prasanth et al. 2005) was found in mouse liver cells. The *mCAT2* protein is a cell-surface receptor involved in the cellular uptake of L-arginine, which is a precursor for the synthesis of nitric oxide (NO). The NO pathway is induced in cells under various stress conditions, including viral infection and wound healing, as a part of cellular defense mechanism. CTN-RNA is transcribed from the same gene as its protein-coding counterpart (mCAT2) but from an alternative promoter, also poly(A) site is different, which results in unique untranslated regions - 5 ‘UTR and 3’ UTR. Upon stress, CTN-RNA is post-transcriptionally cleaved in its 3’UTR, and the released mRNA, similar to mCAT2 mRNA, is transported to the cytoplasm and translated into mCAT2 protein. This mechanism provides a rapid response, in the form of mCAT2 protein production, to stress conditions, which allows modulation of L-arginine uptake for the NO pathway.

Still, there is little information about which mechanisms lead to the retention of nuclear mRNAs, and what causes their release into the cytoplasm. In the case of retention of pre-mRNA, participation of splicing factors was implicated, especially those forming early spliceosome complex (E), i.e. U1 snRNA and U2AF (Takemura et al. 2011). Binding of these factors to mRNAs inhibited export through nuclear pores (Fasken and Corbett 2009). There is also a hypothesis that there are nuclear retention factors, that bind the mRNAs and block the possibility of binding the export factors. It was also expected that these factors anchored mRNAs in nuclear structures (Takemura et al. 2011).

So far, nuclear speckles were the only domain found to be involved in accumulation of polyadenylated transcripts in animal cells. However, the transcripts accumulated in the speckles do not appear to be transported to the cytoplasm (Huang et al. 1994). In larch microsporocytes during diplotene, lasting about 5 months in this species, large quantities of polyadenylated RNA are synthesized (Kołowerzo-Lubnau et al. 2015). We observed that in mid-diplotene poly(A) RNA was accumulated in spherical structures within the nucleoplasm. We demonstrated, using *in situ* hybridization at the ultrastructural level as well as double labelling of poly(A) RNA and known markers of plant Cajal bodies (CBs), that CBs were sites of poly(A) RNA accumulation (Kołowerzo et al. 2009). We also showed that the transcripts accumulated in CBs were indeed encoding proteins - i.e. they were mRNAs. These included housekeeping gene transcripts, e.g. mRNAs encoding pectin methyltransferase, ATPase, catalase, peroxidase, or α-tubulin. In addition to these transcripts, we also localized transcripts encoding proteins commonly found in CBs, e.g. splicing factors SmD1, SmD2, SmE as well as mRNAs encoding RNA polymerase II subunits - RPB2 and RPB10 (Smoliński and Kołowerzo 2012).

CBs are evolutionarily conserved structures occurring in both animal (Morris 2008) and plant cells (Shaw and Brown 2004), which indicates their fundamental role in eukaryotic cells. Intensive research, carried out for many years, has demonstrated the relationship between CBs and many processes associated with the metabolism of various types of RNA, mainly snRNA (Frey and Matera 2002, Kiss et al. 2003, Stanek et al. 2006) and snoRNA (Lamond et al. 2003). CBs are self-organized structures (Mistelli 2007), appearing when there is a need for increased metabolism of certain types of RNA. Although assembly of spliceosomal subunits is possible in the nucleoplasm, snRNPs assembly rate in CBs was found to be around 10-fold greater than in the former (Klingauf et al. 2006, Novotny et al. 2011). In contrast, fibroblasts, cells with low metabolism that do not require intensive assembly of snRNPs, do not have CBs. Here, snRNPs assembly takes place without CBs and therefore it is many times less efficient. Coilin gene (encoding a marker protein for these bodies) knockdown in cells with high metabolism (generative cells or embryonic cells) caused the breakdown of CBs and led to death of these cells (Strzelecka et al. 2010).

Genes in larch microsporocytes were showed to be transcribed in bursts. Five such bursts were observed during diplotene. We demonstrated that a large quantity of poly(A) RNA in the nuclei of microsporocytes appeared at the launch of transcription. These transcripts were retained in the nucleus for a long time and were stored within the nucleoplasm and CBs. Pool of transcripts within the CBs gradually increased and reached a maximum at the stadium preceding the release of large quantities of poly(A) RNAs to the cytoplasm. Cajal bodies are the compartment in which polyadenylated transcripts are retained for a very long period of time (of the order of days). Nuclear pool of these transcripts was stored mainly in Cajal bodies during the stages ending poly(A) RNAs flow cycle (Kołowerzo-Lubnau et al. 2015). Therefore it seems that CBs exert a significant effect on the nuclear retention of mRNA, as well as on the subsequent export of these transcripts to the cytoplasm.

For further detailed analysis, we selected transcripts encoding Sm proteins, which, as we have shown in previous studies, retain in the nucleoplasm and CBs (Smoliński and Kołowerzo 2012).

To examine which mRNAs after nuclear retention are transported to the cytoplasm and potentially are translated, we performed transcriptomic analysis of the cytoplasmic mRNAs. Among these transcripts, mRNAs encoding proteins associated with the process of transcription and post-transcriptional modifications constituted a large group including mRNA encoding Sm proteins. We examined their distribution during a single cycle of cellular synthesis and turnover of poly(A) RNA, and checked in what form they retain in the nucleus. We also checked whether these transcripts are released into the cytoplasm after the retention period and whether the increase in these mRNAs in the cytoplasm correlates with the increase in the level of proteins they encode, which would suggest that nuclear retention is an element of regulation of expression of these mRNAs.

## Results

Our previous studies showed that in larch microsporocytes, five bursts of *de novo* transcription during diplotene corresponded to five cycles of increased levels of poly(A) RNAs in the cell (Kołowerzo-Lubnau et al. 2015). Interestingly, in the mid-diplotene, the colocalization of nuclear poly(A) RNAs with newly formed transcripts was relatively low (21% of total poly(A) RNAs pool). This suggested the occurrence of nuclear poly(A) RNA retention during this period. The observation led us to perform further, detailed analysis of mRNA metabolism in the mid-diplotene.

To address the issue of RNAs nuclear retention, we performed spatial and quantitative analysis of a single cycle of cellular synthesis and turnover of poly(A) RNAs. The characteristic changes in poly (A) RNAs distribution allowed us to distinguish five stages in this cycle. Due to its extraordinarily long duration in our model cells, it was possible to observe and analyze sequence of stages of poly(A) RNAs life cycle. At the beginning of the cycle, during the first two stages, amount of nuclear poly(A) RNAs was **significantly** higher than in the cytoplasm (Figures 1 and 2A). Nuclear pool of poly(A) RNAs was in part dispersed, but also localized within numerous spherical structures corresponding to Cajal bodies (CBs) (Figure 1). At the initial stage, the level of poly(A) RNAs was the highest (Figure 2A) and was correlated with high level of hyperphosphorylated form of RNA polymerase II (Pol II O) staining (Figure 1). At this stage the burst of polyadenylated RNA synthesis *de novo* occurred. Starting from the second stage, the nuclear pool of poly(A) RNAs progressively decreased, with concomitant increase in the cytoplasmic pool (Figures 1 and 2A). During this stage we still observed a pattern of distribution similar to the one described above (Figure 1), however the proportion of poly(A) RNAs in cytoplasm and CBs was changed. The abundance of poly(A) RNAs in CBs constantly increased throughout the cycle (Figure 2B), and along with the rearrangement of poly(A) RNAs nuclear distribution, gradual increase of their abundance in the cytoplasm occurred (Figure 2A). Poly(A) RNAs accumulated within the perinuclear region in a characteristic pattern, which we called “ER-like” (Figure 1). During the third stage a dramatic drop in nucleoplasmic pool of poly(A) RNAs was noted, which was accompanied by increased staining of the cytoplasm (Figure 2A), where it was more dispersed compared to the previous stage (Figure 1). Poly(A) RNAs signal decreased also in CBs (Figure 2A), although the ratio of these transcripts within CBs versus total nuclear pool was only up to 5% higher than in the second stage, and up to 22% higher than in the first one (Figure 2B). On the microscopic level we could observe numerous strongly stained CBs at this stage (Figures 1). The fourth stage was characterized by further decrease of nucleoplasmic level of poly(A) RNAs (Figure 2A), whereas a significant portion (41% of nuclear pool) of transcripts still accumulated in numerous CBs (Figures 1 and 2B). This correlated with the increase of poly(A) RNA level within the cytoplasm (Figure 2A). Numerous bright foci were observed, scattered over the whole cytoplasmic area (Figure 1).

**Figure 1.**
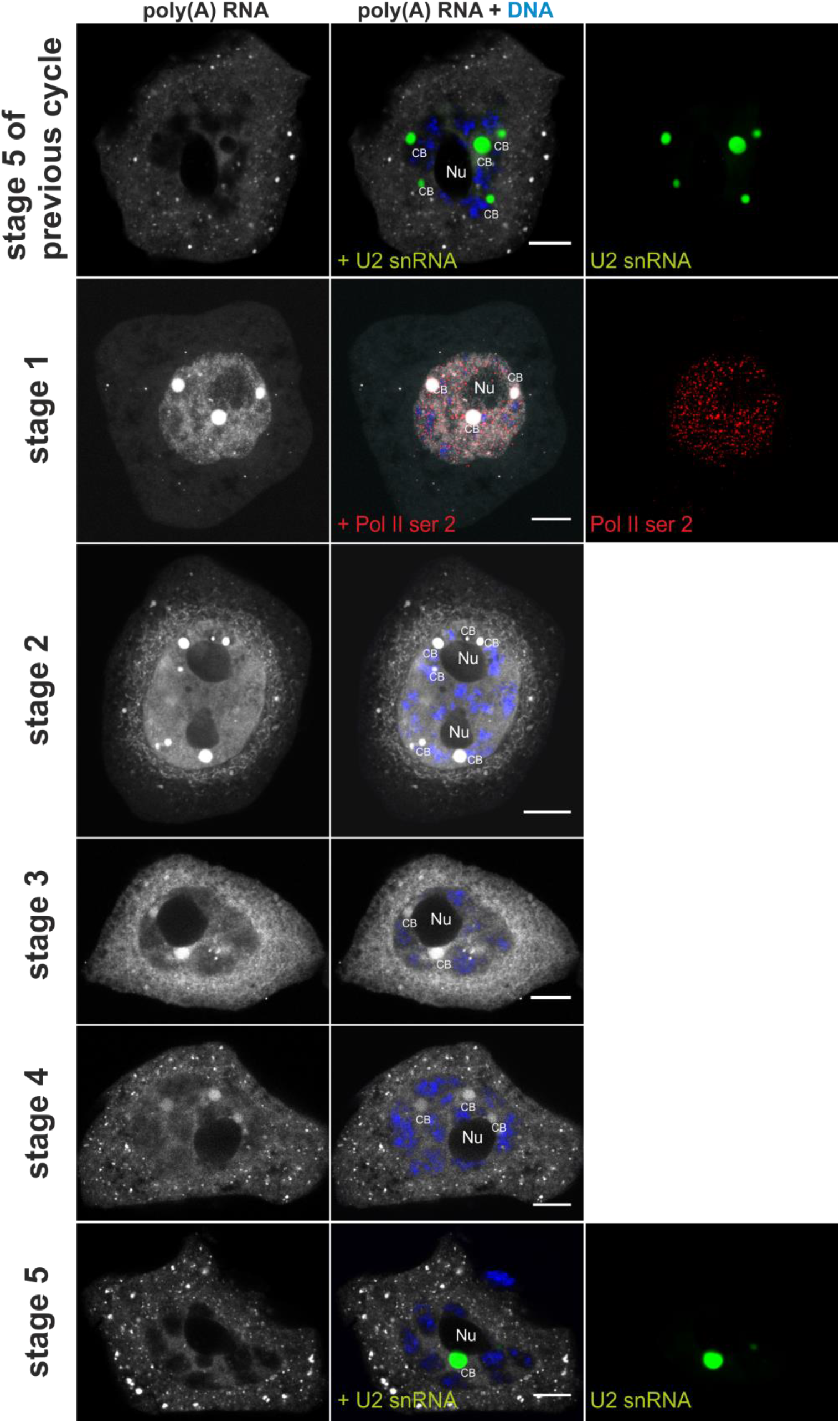
The cycle of cellular synthesis and turnover of poly(A) RNA. U2 snRNA. Detailed description is provided in the text; Labeling of U2 snRNAs (green) was used to show CBs at stage 5 when poly(A) RNAs do not accumulate in them. Labeling of RNA polymerase II (red) in active form (phosphorylation of serine 2 in the CTD domain of the polymerase) was used to indicate the stage with the highest transcriptional activity. CB – Cajal body, Nu – nucleolus. Bar - 10μm.

**Figure 2.**
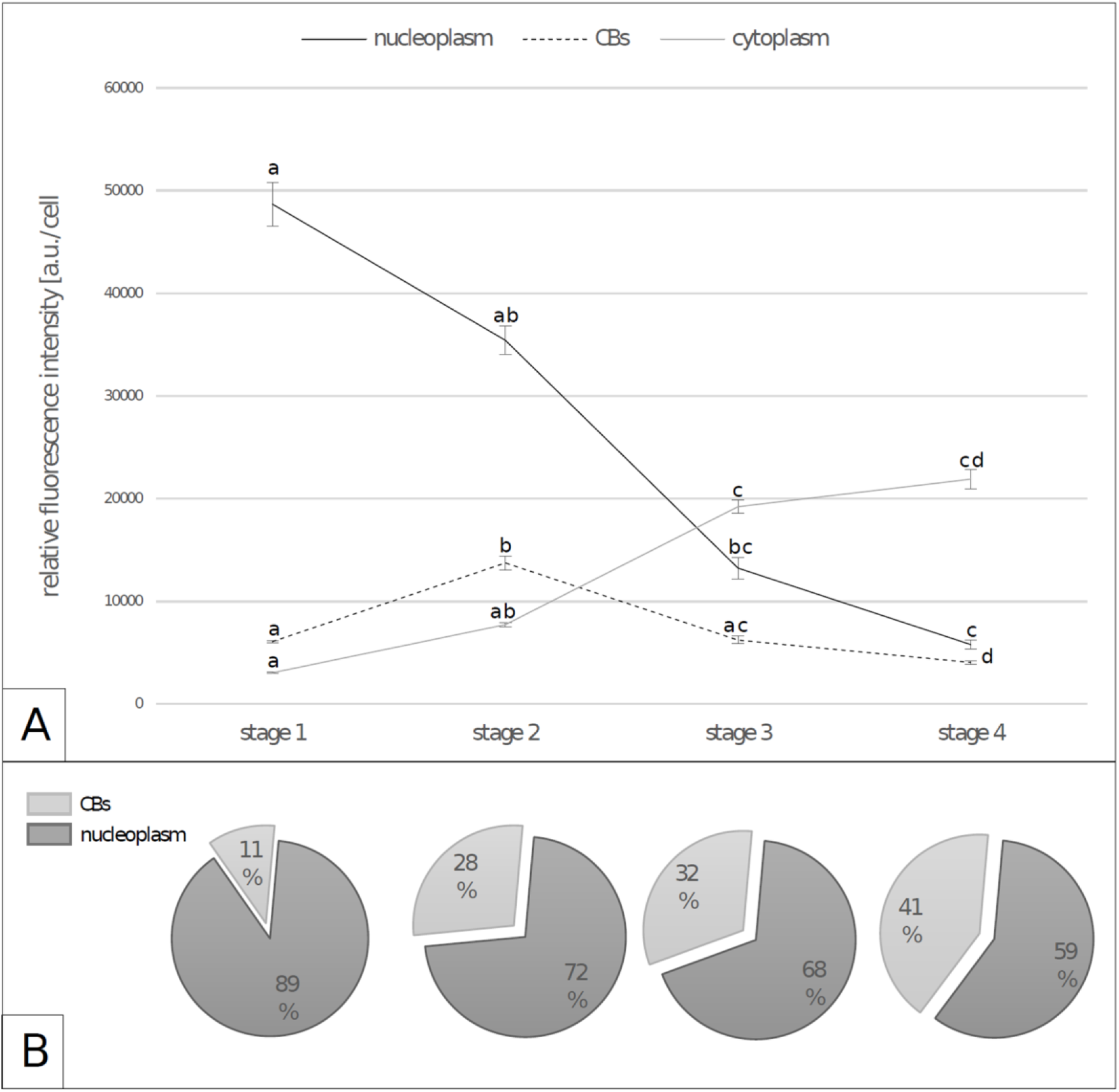
The level of poly(A) RNA in cellular compartments during the cycle of cellular synthesis and turnover of poly(A) RNA. **A.** The level of poly (A) RNA in individual cell compartments (nucleoplasm, CBs, cytoplasm) during the single cycle. Whiskers indicate Standard Error of the Mean (SEM). Significance of differences between stadia (at a=0.05), according to Kruskal-Wallis test with Dunn’s post-hoc procedure using Benjamini-Hochberg FDR correction, is denoted with different letters (i.e. the same letter tells there is no difference). **B.** Percentage of the nucleoplasmic and CBs poly (A) RNA fraction in the nucleus during a single cycle.

Fifth stage was short-lived, being quick transition between successive cycles and we were able to observe only few cells. At this stage, the level of poly (A) RNA in nucleoplasm was the lowest in the whole cycle. There was also no accumulation of these RNAs in CBs (Figure 1). Due to the very small number of cells at this stage it had not been subjected to quantitative analysis.

The presented data indicated that the phenomenon of temporal nuclear retention of RNA occured in larch microsporocytes. The high nuclear level of poly(A) RNAs continued throughout the majority of the cycle, long after *de novo* transcription was silenced. These results confirm that a significant portion of newly transcribed poly(A) RNAs was not immediately subject to cytoplasmic export, but was temporarily retained within the nucleus. Interestingly, the nuclear domains involved in this process were most likely CBs. Despite the fact that starting from the second stage the nuclear pool of RNA continuously decreased, the ratio of transcripts abundance within CBs versus total nuclear pool had increased, which points at CBs as the sites of RNA nuclear retention. The concentration of poly (A) RNAs in CBs was 15 to 70 times higher than in the rest of the nucleus, despite the fact that the volume of Cajal bodies constituted only 0.5% −1.5% of the volume of the nucleus. These small structures accumulated up to 40% of the total pool of poly (A) RNAs in the nucleus.

In recent years, there were several publications about nuclear retention in both animal and plant cells. According to these works, transcripts were retained in various forms of maturity. The main conclusion of these studies was that the nuclear retention of mRNAs aims to control the time of their export to the cytoplasm and thereby the time of translation.

To get information on which mRNAs were transported to the cytoplasm and were potentially translated during a single cycle of cellular synthesis and turnover of poly(A) RNA we performed the analysis of the cytoplasmic transcriptome of larch microsporocytes. As a result of this analysis, we received 7 001 transcript sequences located in the cytoplasm (Figure 3A). To identify these transcripts, the sequences were analysed using BLASTX (for potential mRNA sequences) and BLASTN (for other sequences) (NCBI database for green plants was searched). After identifying the coding sequences, a functional annotation was made. 3,283 sequences (44%) were assigned a total of 307 714 GO terms (Gene Ontology). The cytoplasmic transcripts of larch microsporocytes are listed in Supplementary Data 1. Functional analysis showed that cytoplasmic transcripts included 5 major mRNAs groups encoding proteins associated with: (1) transcription and posttranscriptional modification processes, (2) ribosomes, translational and post-translational modification of proteins, (3) mitochondria and energy transformations, (4) photosynthesis and plastids, (5) a cytoskeleton and a cell wall (Figure 3 B). The most numerous group were transcripts encoding proteins associated with transcription and post-transcriptional modifications, among which were transcripts coding for Sm proteins: D1, D2, and G. Sm are nuclear proteins involved in pre-mRNA splicing by being a core component of spliceosomal ribonucleoproteins (UsnRNPs) and in plants are considered one of CBs markers. From our previous studies we know that these specific mRNAs are retained in the nucleus and are accumulated in the CBs during the diplotene microsporocytes larch (Smoliński & Kołowerzo 2012), which makes these transcripts particularly interesting for our research and these mRNAs we used for further analysis. We made an attempt to assess in what form: mature or immature, the transcripts coding for Sm proteins were retained in the nucleus. As the sequences obtained from the cytoplasmic transcriptome were mature transcripts and the larch genome was not sequenced, we had to try to sequence the genes coding for Sm proteins to get information about immature transcripts. Using the sequences obtained from the larch transcriptome, as well as the genomes of related species, mainly *Picea glauca* and *Picea abies*, we designed primer pairs amplifying intron sequences together with flanking exons fragments. We obtained PCR products that were sequenced and used to design probes for fluorescent in situ hybridization and PCR primers.

**Figure 3.**
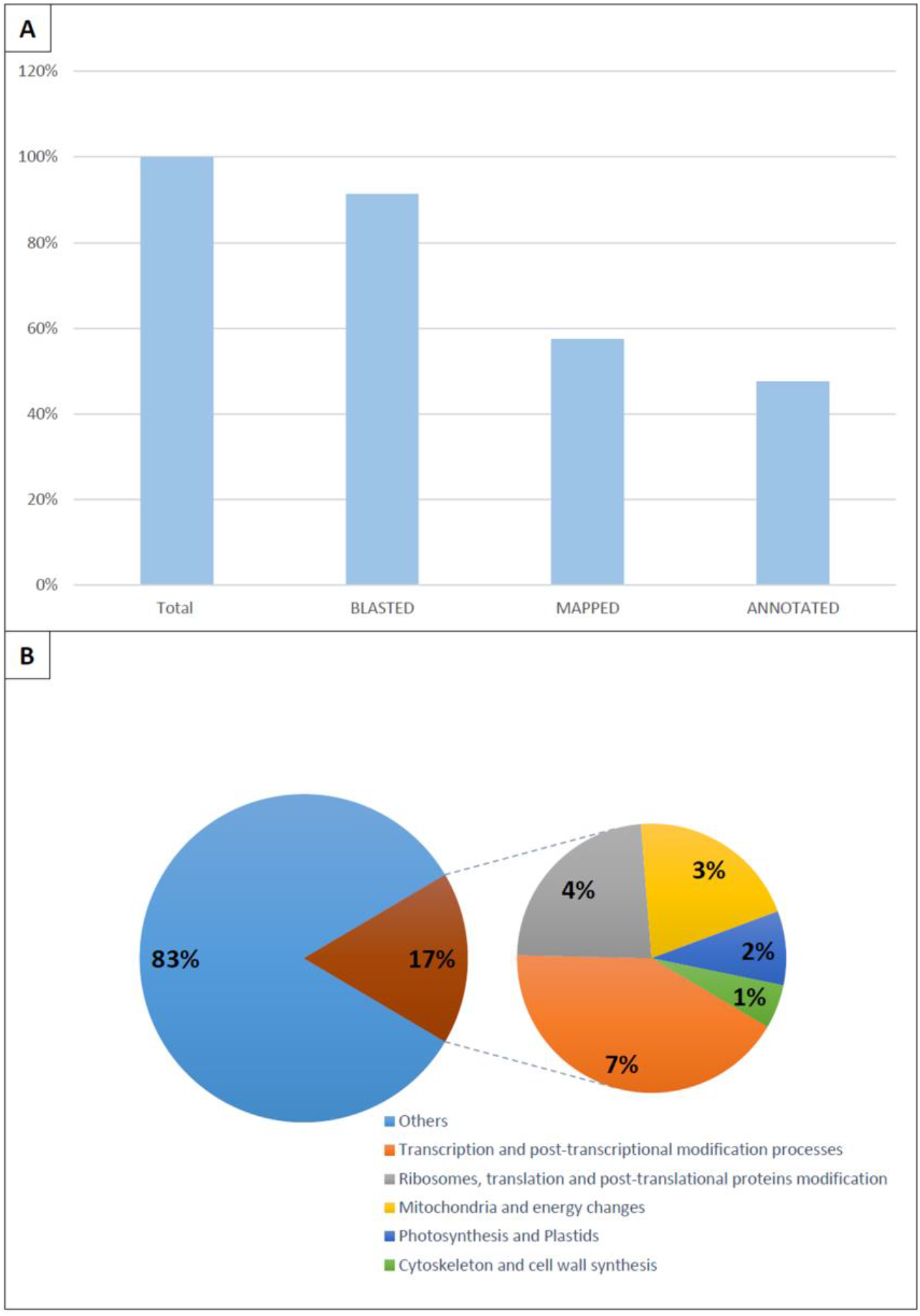
Annotation of cytoplasmic mRNAs. **A.** Distribution of sequences on annotation levels. 100% is 7001 sequences. **B.** Functional annotation of cytoplasmic mRNA at level 3 of biological process domain.

FISH analysis showed that all the analyzed transcripts coding for Sm proteins are retained in the microsporocytes nucleus in immature form (Figure 4). In the case of pre-mRNA for SmD1, consisting of three exons and two introns, the signal coming from the sequence complementary to both the first and second intron was accumulated in nucleoplasm as well as in CBs (Figure 4A). On the other hand, in the case of pre-mRNA coding SmD2 and SmG proteins, which consisted of four exons and three introns, the signal from only one intron showed accumulation in nucleoplasm and CBs (Figures 4B and 4C). The signal from the other two introns was either very weak or limited only to a very small area in the nucleus which may indicate that they are quickly spliced from the transcripts. These transcripts containing introns exhibit nuclear accumulation. Unspliced introns, so-called retained introns (IR), determines the retention of such a transcript in the nucleus.

**Figure 4.**
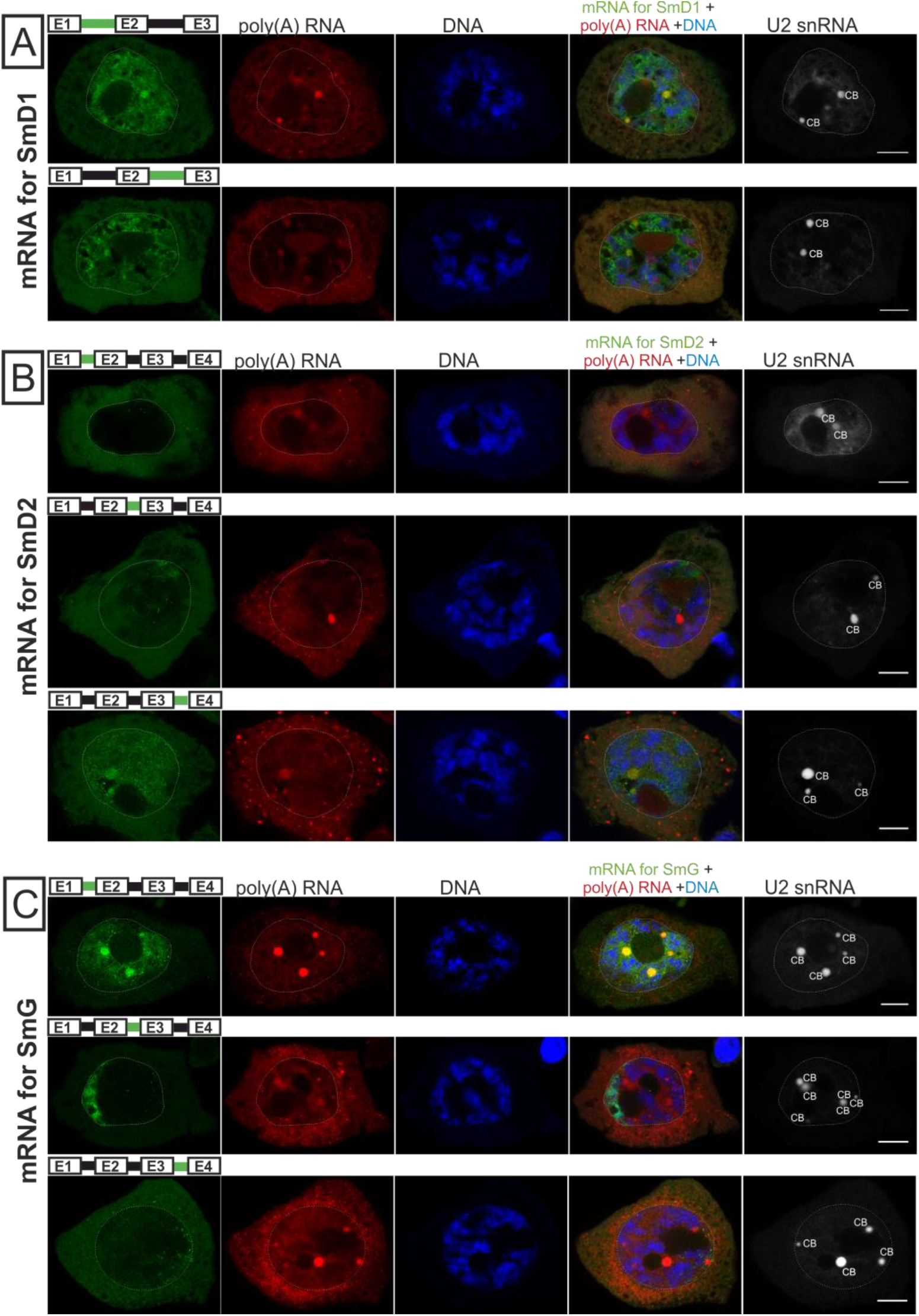
mRNAs coding for Sm proteins are retained in nucleoplasm and Cajal bodies (CBs) as pre-mRNAs containing defined intron(s). **A.** Localization of pre-mRNA encoding the SmD1 protein. Labeling of the intron sequences present in the mRNA (green) showed that both the transcript with the first (upper panel) and second intron (lower panel) accumulate in the nucleoplasm and Cbs, which was confirmed by co-localization with U2 snRNA (fifth column). **B.** Localization of pre-mRNA encoding the SmD2 protein. Labeling of the intron sequences present in the mRNA (green) showed that only the sequence containing the third intron is accumulated in the nucleoplasm and CBs (lower panel), while the sequences containing the first (higher panel) and second intron (middle panel) are visible only in a few clusters. **C.** Localization of pre-mRNA encoding the SmG protein. Labeling of the intron sequences present in the mRNA (green) showed that only the sequence containing the first intron is accumulated in the nucleoplasm and CBs (higher panel) while the sequences containing the second (middle panel) and third intron (lower panel), occurs only on the periphery of the nucleus. CB – Cajal body. Bar - 10μ.

In further studies, we conducted a more detailed analysis of the transcript encoding the SmG protein. In this case, the pre-mRNA consists of four exons and three introns, and our studies show that the first intron is a retained intron. Using multi-FISH labelling, we analyzed the location and level of pre-mRNA coding for SmG protein with the first retained intron in the single cycle of cellular synthesis and turnover of poly(A) RNA (Figure 5). These transcripts were accumulated in both CBs and nucleoplasm up to the fourth stage of the cycle (Figure 5A). In the fifth stage, the level of labeling was very low, we only observed point signals from CBs and nucleoplasm. Quantitative analysis revealed that the level of the pre-mRNA gradually decrease both CBs and nucleoplasm from the second cycle stage (Figure 5B). Control reactions have shown that, in fact, retained mRNAs encoding SmG proteins only occur with the first intron retained, while the second intron sequence is not retained, which indicates selectively retaining a particular intron (Figure 5C).

**Figure 5.**
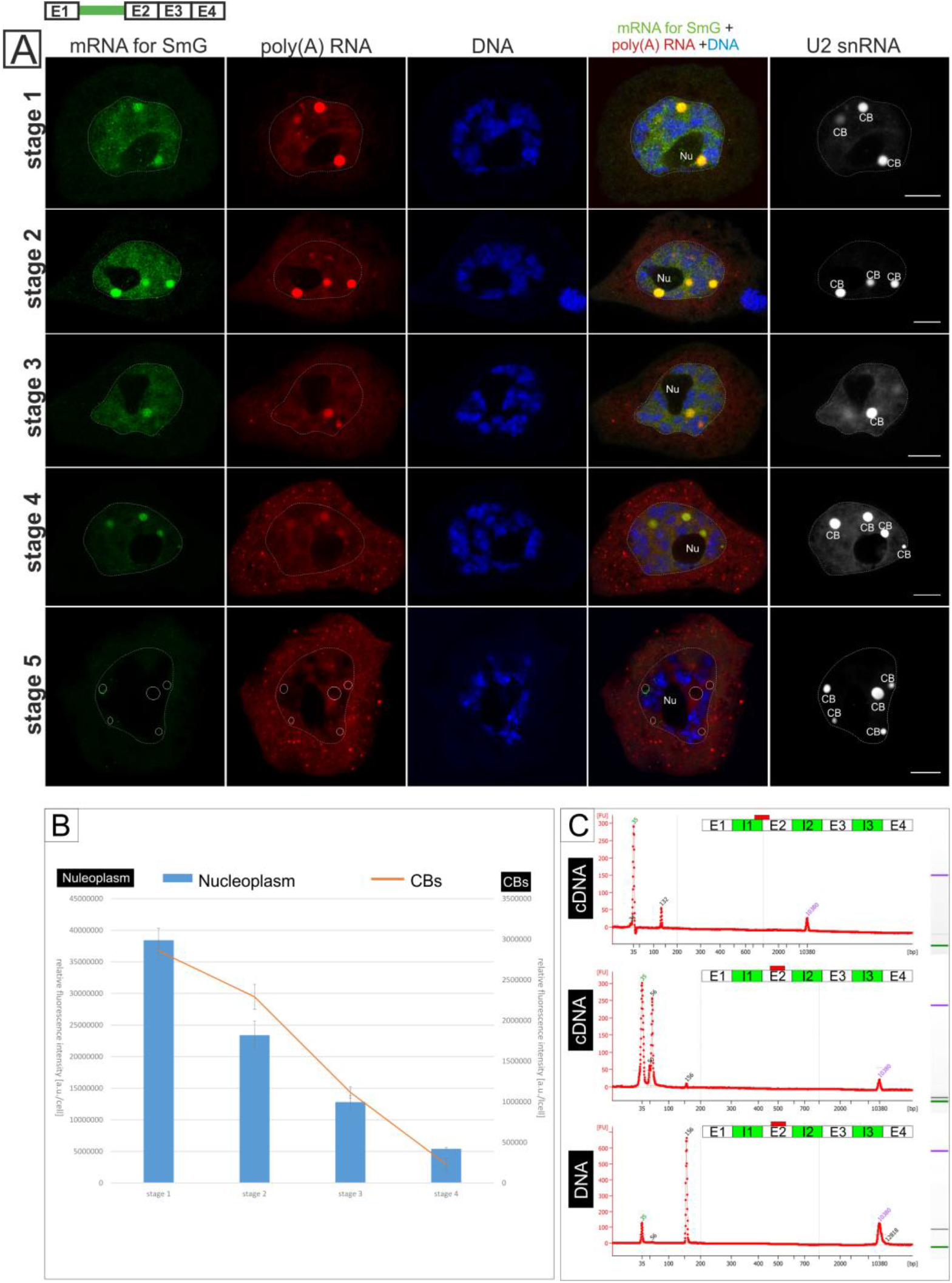
Detailed analysis of pre-mRNA coding for SmG proteins with the first retained intron. **A.** Distribution of the pre-mRNA encoding the SmG protein with the first intron sequence unspliced against the background of the general pool of poly(A) RNA distribution. U2 snRNA staining was used to show Cajal bodies (CB). Details in the text. Nu – nucleolus, CB – Cajal body. Bar - 10μ. **B.** Level of pre-mRNA coding for SmG proteins with the first retained intron in the nucleoplasm (upper panel) and CBs (lower panel). Whiskers indicate Standard Error of the Mean (SEM). Significance of differences between stadia (at a=0.05), according to Kruskal-Wallis test with Dunn’s post-hoc procedure using Benjamini-Hochberg FDR correction, is denoted with different letters (i.e. the same letter tells there is no difference).**C.** High Sensitivity DNA Bioanalyzer assay as a control for selective retention of particular introns. Intron 1 sequence can be PCR-amplified (132 bp, upper panel) from cDNA. In contrast, no product for intron 2 (156 bp) can be generated on cDNA (middle panel), but it is amplified on DNA (lower panel).

To check whether and when retained introns are spliced with pre-mRNA we designed a probe for the fully mature mRNA after splicing the first intron and checked the location of such a transcripts during a single cycle of cellular synthesis and turnover of poly(A) RNA (Figure 6). We have already observed the fully spliced transcript in the first stage. In first stage it was present mainly in nucleoplasm we did not observe its accumulation in CBs. Probably splicing of the retained introns takes place immediately after the release of such a transcripts from CBs and before its export to the cytoplasm. In subsequent stages, fully spliced transcripts were present both in the nucleus and in the cytoplasm. In the second stage, even nuclear and cytoplasmic labeling was observed, while in subsequent stages, the nuclear labeling drastically decreased in favor of the cytoplasmic signal. Probably the rate of export and further use of these transcripts vary depending on the stage of the cycle. In the first two stages, the significant accumulation of fully mature transcripts in the nucleus may suggest that the rate of their export to the cytoplasm is not too high. However, in subsequent stages, the nuclear signal decreases significantly and there is more of it in the cytoplasm, which may mean a significant increase in the export rate.

**Figure 6.**
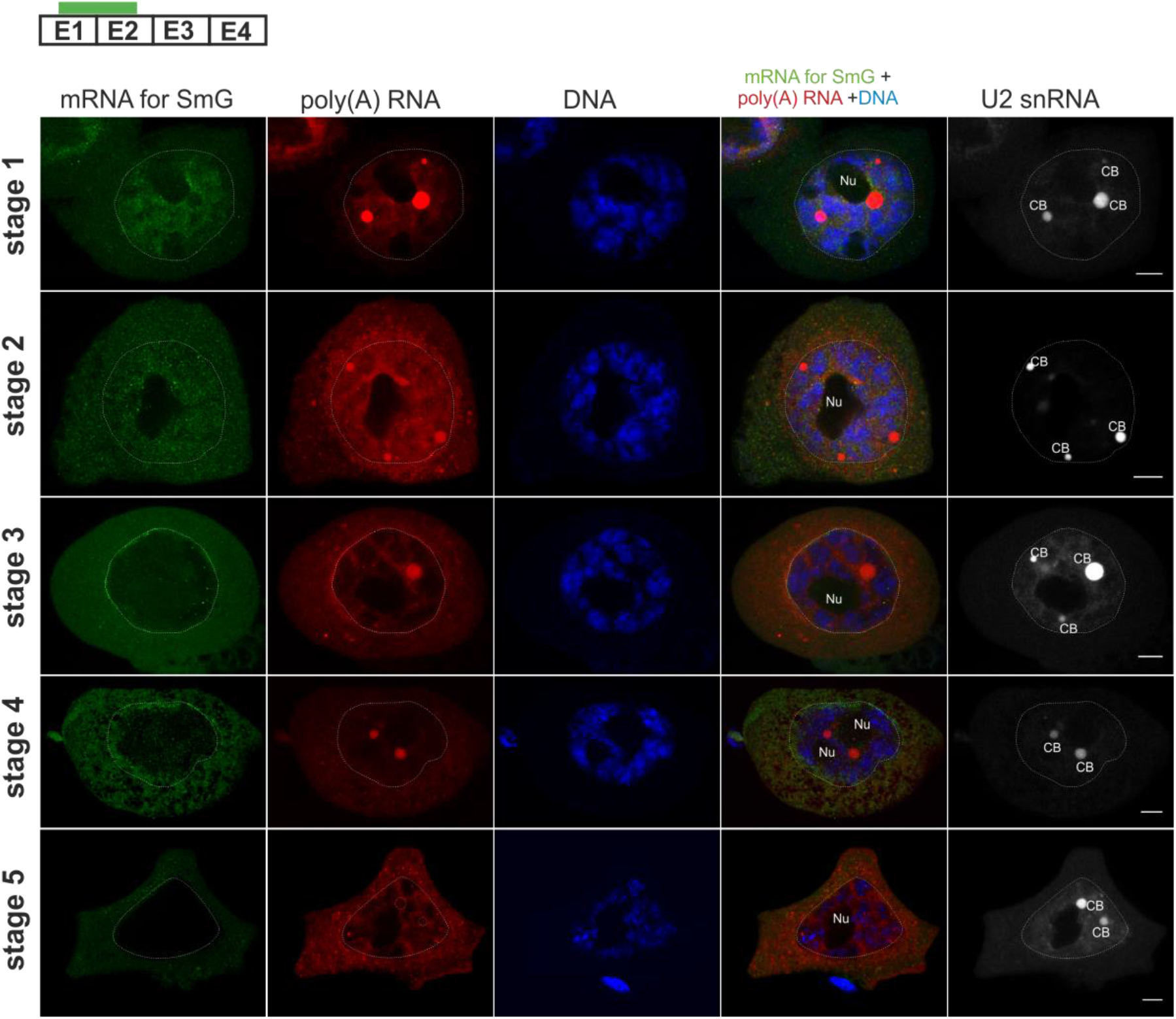
After removal of the retained intron, the mRNAs are no longer accumulated in CBs. Mature SmG-encoding mRNA is absent from CBs during all stages. During stages 1 and 2 it is present in the nucleoplasm, concentrates on nuclear membrane in stage 3, while during stages 4 and 5 it is present in cytoplasm. Bar - 10μ.

Detailed analysis of the transcripts encoding SmG proteins showed a clear involvement of CBs in the retention of these mRNAs in the cell nucleus. As research has shown, only the non-fully spliced form of transcripts with retained introns are present in CBs, while the mature form, which are exported to the cytoplasm, are not accumulated. A question arises whether the mRNAs retained in the nucleus maintain their function, i.e. whether they are translated after released to the cytoplasm. In order to address this question, we analyzed a single cycle of expression of a specific protein coding gene, with emphasis on the relation between mRNAs retention in CBs and the latter protein synthesis in the cytoplasm. For the analysis, a gene family coding for Sm proteins was selected, namely SmD1, SmD2 and SmE. We conducted a quantitative analysis of the level of mRNAs encoding Sm proteins during a single cycle in individual microsporocyte compartments: in nucleoplasm, Cajal bodies and cytoplasm. We observed a special correlation between the increase in mRNAs signal in CBs and the appearance of mRNAs in the cytoplasm. An increase in mRNAs level in CBs is manifested by the appearance / increase of its cytoplasmic pool in the next stage (Fig 7A).

**Figure 7.**
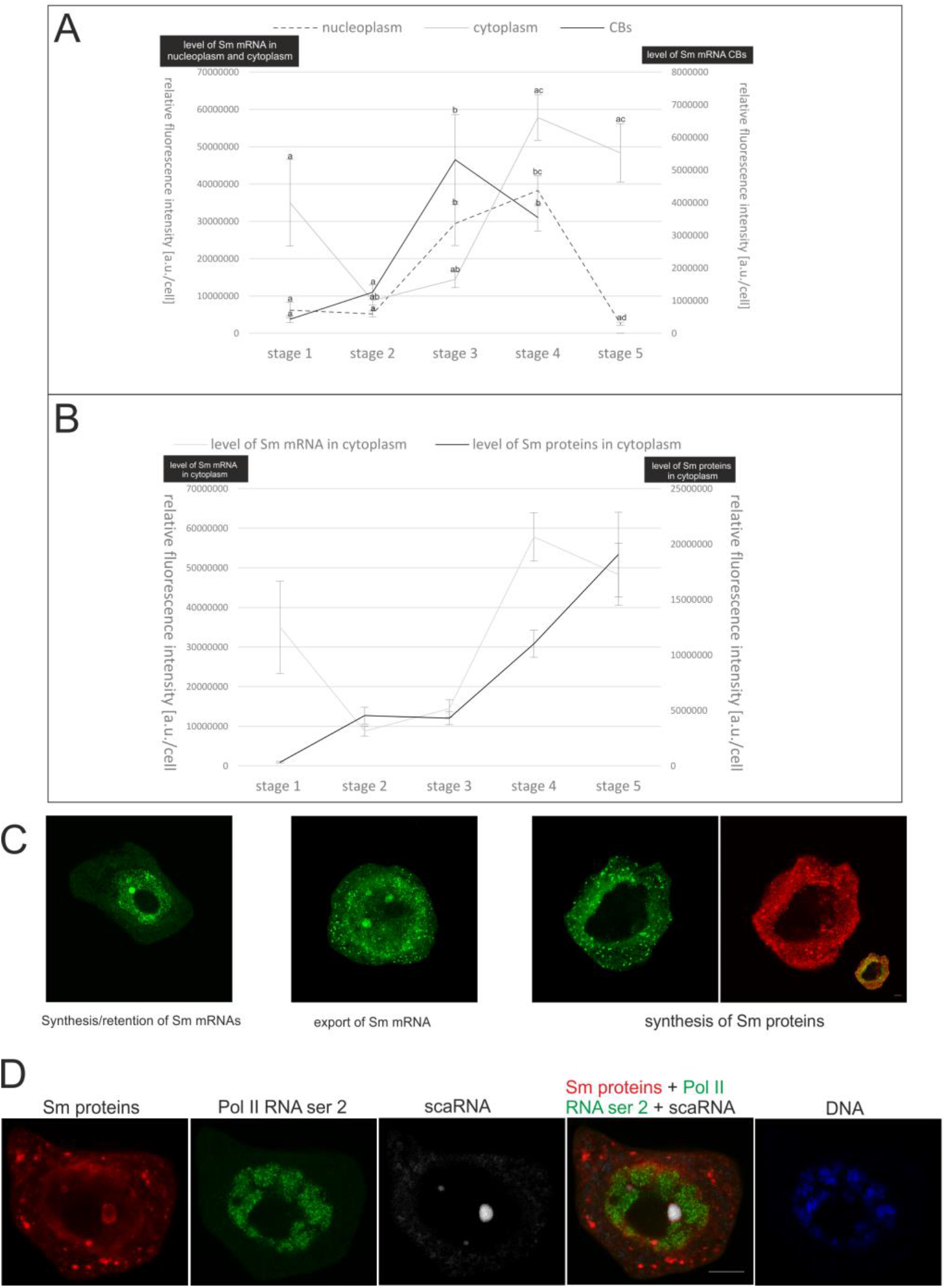
Retained mRNAs are exported to the cytoplasm and translated. **A.** Level of mRNA encoding Sm proteins during a single cycle. **B.** Comparison of the cytoplasmic pool of mRNA encoding Sm proteins to the cytoplasmic pool of Sm proteins during a single cycle. The two variables are positively correlated (Spearman’s rank correlation, r=0.56, p=0, n=55). **C.** Visualization of Sm protein-encoding genes expression stages. **D.** First stage of the next cycle of poly(A) RNA synthesis and turnover. High level of pol II RNA phospho CTD Ser 2 indicates the start of another transcriptional burst, while new Sm proteins are formed in the cytoplasm, from the mRNA pool that was processed in the previous cycle. scaRNA was used as a marker of CBs. CB-Cajal body. Bar 10μ.

This condition persists until the third stage, in which the mRNA encoding Sm proteins reaches their maximum level in CBs. In subsequent stages, this level decreases, and the level of mRNAs encoding Sm proteins in the cytoplasm remains very high, which may indicate intensive translation.

To confirm that the mRNA exported to the cytoplasm area was translated, we also examined the level of Sm proteins with particular emphasis on the cytoplasm - the place where we could observe the formation of a new pool of proteins. As we thought, the level of Sm proteins in the cytoplasm increased rapidly in the last stages of the cycle, when we observed a rapid decrease in the mRNAs pool in CBs and a high level of mRNAs in the cytoplasm (Figure 7B). This is also confirmed by microscopic observations. In stage five, where in microsporocytes mRNA encoding Sm proteins are observed almost exclusively in the cytoplasm, we also observe intensive protein labeling only in this compartment, probably reflecting *de novo* Sm protein synthesis (Figure 7C). It can therefore be assumed that the retention of the not fully spliced form of mRNAs in larch microsporocytes is of a functional nature serving to regulate the time of export and thus regulating the time of synthesis of individual proteins. It seems that this Sm protein synthesis at the end of the cycle is intended to provide splicing factors that will be necessary at the beginning of the next cycle, where transcription activity resumes (Figure 7D).

## Discusion

### Nuclear retention of mRNA in larch microsporocytes is a mechanism controlling export and translation time

As has been found in many other species, including mammals (Halpern et al. 2015), transcription during prophase in larch microsporocytes, is a pulsating process (Kołowerzo-Lubnau et al. 2015). As our previous studies have shown, each transcription burst generates a poly (A) RNA pool which is retained for some time in the cell nucleus after the transcriptional activity of microsporocytes is silenced. This is related to the rearrangement of chromatin during meiotic prophase. In larch microsporocytes, during diplotene, there are four phases of chromatin contraction and 5 diffusion phases, when its relaxation occurs (the so-called diplotene diffusion stage) (Kołowerzo-Lubnau et al 2015). Many studies indicate that the retention of mRNA in the cell nucleus is the result of mechanisms of quality control and detection of defective mRNAs and their subsequent degradation (Fasken and Corbett 2009). In larch microsporocytes, the level of poly (A) RNAs that are retained within the nucleus is very high. Their nuclear pool significantly exceeds the cytoplasmic one throughout the entire period of diplotene (Kołowerzo-Lubnau et al. 2015). The scale of this phenomenon suggests that the retention of nuclear mRNA in larch microsporocytes plays a new, unappreciated role in the expression regulation of certain genes, rather than just capturing defective transcripts. Our research showed that mRNAs retained at the beginning of the cycle of cellular synthesis and turnover of poly(A) RNA were exported to the cytoplasm, initially in lower and then higher quantity. We assumed that the export of mRNAs to the cytoplasm will be associated with their translation and the biosynthesis of functional proteins. To test this hypothesis, we analyzed the cytoplasmic transcriptome of larch microsporocytes to identify mRNAs present at stages that are characterized by high export and high levels of mRNA in cytoplasm. In the analyzed cytoplasmic transcriptome, the most numerous group were transcripts encoding proteins associated with transcription and post-transcriptional modifications, among which were transcripts coding for Sm proteins: D1, D2, and G. Furthermore, our previous research showed that Sm proteins and other splicing factors are intensely synthesized for the needs of processing products of transcription burst occuring in diplotene (Hyjek et al. 2015). Therefore, we chose mRNAs encoding Sm proteins for further analysis. We checked whether the level of exported mRNAs coding for Sm proteins correlates with the level of newly synthesized Sm proteins. We confirmed that at the last stage of the cycle in which mRNAs coding for Sm proteins were almost exclusively in the cytoplasm, we also observed newly synthesized Sm proteins. This indicates that we are dealing here with the synthesis and retention of functional transcripts, which are exported and translated over time. Moreover, it points out that nuclear retention of mRNAs encoding Sm proteins is aimed at introducing a new pool of Sm proteins at a very specific moment in a cycle. Earlier studies on the biosynthesis of snRNPs complexes in larch microsporocytes have shown that Sm proteins and snRNAs are synthesized de novo in pulses (Smoliński et al. 2011, Hyjek et al. 2015). Thus, the assembly of functional snRNPs complexes takes place in a cyclical manner, as individual elements of these complexes become available in a cell. We have shown that the moment of appearance of newly synthesized Sm proteins coincides with the start of a new transcription burst. This indicates that the retention of mRNAs encoding Sm proteins in the nucleus delays the translation time of these proteins to the next transcriptional round. This means that when the cell begins its high transcriptional activity again, it will already be equipped with elements of snRNPs complexes necessary in the splicing process. The appearance of Sm proteins in the microsporocyte cytoplasm initiates the assembly of functional snRNPs complexes. It is a multistage process, including stages such as transcription of snRNAs, export of pre-snRNAs into the cytoplasm, Sm core assembly and exonucleolytic trimming of the snRNAs’ 3′ ends, hypermethylation of 7-methylguanosine (m7G) cap to form a 2,2,7-trimethylguanosine and re-import of snRNPs into the nucleus, where they are directed to CBs for nuclear maturation steps (Matera and Wang 2014, Bohnsack and Sloan 2018). The complexity of this process probably means that access to fully functional snRNPs complexes at the time of transcription burst in microsporocytes is likely to be limited. This deficit of fully functional splicing complexes, in addition to the mechanism of self-regulation of the level of splicing proteins, such as Sm proteins, may complement the retention mechanism of mRNAs generated in a single transcriptional pulse. Insufficient splicing factors can affect the selection of pre-mRNAs that will be spliced first and which will be retained for later processing. A specific feedback mechanism is created, depending on the availability of splicing factors. Naro et al. (2017) showed that the direct cause of intron retention is the low efficiency of their splicing, related to the high activity of RNA polymerase II. The high intensity of transcription causes competition between individual introns for the attachment of splicing factors, which in turn leads to the retention of introns with weak splicing sites, usually characteristic of retained introns. It has been shown that decreasing the intensity of transcription by inhibiting the phosphorylation of RNA polymerase II, results in a significant improvement in the splicing efficiency of retained introns, which indicates that these introns bind spliceosomes less well, and therefore lose the competition with other introns, and therefore remain unspliced.

Mechanisms of functional nuclear retention have been described, among others, in mammalian cells (Halpern et al. 2015). It has been shown that a wide range of genes in highly metabolically active tissues, generates fully spliced, polyadenylated transcripts that are retained in the nucleus for periods that exceed their cytoplasmic lifetimes. These include genes such as the transcription factor ChREBP, Nlrp6, glucokinase, and glucagon receptor. This study demonstrated that nuclear retention of mRNA could efficiently buffer cytoplasmic transcript levels from noise that arises from transcriptional bursts.

An interesting example of functional retention is the formation of an mRNA reservoir in the nucleus of cells that will be transcriptionally silenced for an extended period of time. In the microspores of fern *Marsilea vestita*, it has been shown that despite completely silenced transcriptional activity, at certain times in the development of these cells, proteins necessary for the progress of spermatogenesis are synthesized (Boothby et.al 2013). Analysis of RNAseq-derived transcriptomes revealed a large subset of intron-retaining transcripts that encode proteins essential for gamete development. These transcripts are retained within the cell nucleus, and then at the precise moment of development they are post-transcriptionally spliced, exported to the cytoplasm and translated. These results indicate that the retention of introns arrests translational events necessary for spermatogenesis in *M. vestita*. In male generative cell lines in mice there is a widespread intron retention program in meiosis, which stabilizes highly expressed transcripts with strong relevance for spermatogenesis. Intron-retaining mRNAs are accumulated in the nucleus of meiotic cells of *Schizosaccharomyces pombe* (Averbeck et al. 2005) and mouse (Naro et al. 2017) for hours or days and can be recruited onto the ribosomes for translation long after their synthesis. It indicates that nuclear retention of mRNAs can also favor accumulation, storage, and timely usage of specific transcripts during highly organized cell differentiation programs, such as spermatogenesis.

The positive role of nuclear retention of mRNAs has also been demonstrated in many cells under stress. A known cell response to heat shock is inhibition of splicing, which results in nuclear retention of some pre-mRNAs (Sadis et al. 1988, Saavedra et al. 1996, Shalgi et al. 2014). This mechanism may prevent production of aberrant proteins. Presence of intron-retaining transcripts in the cell at substantial levels suggests that they are relatively stable in the nucleus and may have some function, perhaps serving as a reservoir of partially processed transcripts, which can later be spliced and exported to cytoplasm, to rapidly produce protein when stress conditions abate (Shalgi et al. 2014).

### mRNAs coding for Sm proteins that are retained in larch microsporocytes are not fully spliced

Studies have shown that functional nuclear mRNA retention may relate to both fully mature transcripts (Halpern et al. 2015) as well as to those not fully spliced (Boothby et.al 2013, Shalgi et al. 2014, Bergeron et al. 2015, Naro et al. 2017). Here we have shown that in larch microsporocytes, transcripts encoding Sm proteins are retained in the nucleoplasm and CBs as not fully spliced pre-mRNAs, with one or more introns retained. At precisely defined times, retained introns are spliced, and fully mature mRNAs are released into the cytoplasm, where the proteins are produced.

Intron retention (IR) is a conserved form of alternative splicing characterized by the inclusion of intronic sequence in a mature transcript (Intron Retained Transcripts - IRTs). IR has recently been revealed as an independent mechanism of controlling and enhancing the complexity of transcriptome. IR facilitates rapid responses to biological stimuli, is involved in disease pathogenesis, and can generate novel protein isoforms (Vanichkina et al. 2017). As shown by recent animal studies, the fate of intron-retaining transcripts can vary. IRTs can be exported to the cytoplasm to either generate distinct protein isoforms (Gontijo et al. 2010, Hossain et al. 2016) or be degraded by nonsense mediated RNA decay (NMD) (Jaillon et al. 2008, Peccarelli et al. 2014). However, some intron-containing mRNAs, as documented in yeast (Averbeck et al. 2005), plant (Boothby et.al 2013) and animal cells (Boutz et al. 2015, Naro et al. 2017) remain in the nucleus, leading to time-regulated protein production from the transcript. In this case IR does not lead to generation of another protein isoform (Yap et al. 2012; Braunschweig et al. 2014; Boutz et al. 2015; Bresson et al. 2015; Liu et al. 2017).

In larch microsporocytes, intron-containing mRNAs are retained in the nucleus until splicing is completed. As the example of mRNA coding for SmG protein with the retained first intron shows, the rate of post-transcriptional splicing of this intron regulates the rate of export of this mRNA, and thus regulates the time and level of protein synthesis. This type of regulation is mainly observed in developing, specialized or stressed cells, often with reduced or even inhibited transcriptional activity. Retained transcripts here are a long-term deposit that can be used at a specific time to produce a given protein.

In microspores of *M. vestita,* analysis of the transcriptome at various time intervals during the development of these cells showed that a large proportion of the transcripts usually retain one relatively short intron (Boothby et al. 2013). RT-PCR confirmed that IRTs temporally precede their fully spliced isoforms during development. Studies using spliceosome inhibitor Spliceostatin A and the transcriptional inhibitor α-amanitin confirm that the maturation of IRTs to mRNA was spliceosome-dependent but independent of transcription. These studies show that in the microspores of *M. vestita* the retention of specific introns is regulated during desiccation, a time when the spores undergo expansive translation of stored pre-mRNAs. A similar mechanism of nuclear retention of intron-containing mRNAs was also described in mouse meiotic spermatocytes and post-meiotic spermatids (Naro et.al. 2017). Most intron retention events were detected in meiotic spermatocytes, whereas these introns were spliced in post-meiotic cells. This indicates that these transcripts are stable in the nucleus and can be maintained for days after synthesis and that IR regulation timely determines the usage of transcripts when transcription is repressed. These observations suggested that temporal regulation of IR plays an important role in male germ cell differentiation.

Boutz et al. (2015) demonstrated the presence of intron- retaining mRNAs both in cell lines and liver tissue from adult animals. A common set of such transcripts was found among genes expressed in both embryonic stem cells and adult livers, alongside sets of tissue-specific IRTs that are enriched for genes with particular functions related to the cell type in which they occur. Many of the functional gene categories in which IRTs are overrepresented encode products whose physiological concentrations must be tightly controlled. Regulation of intron- retaining mRNAs splicing kinetics may contribute an additional control point for fine-tuning of gene expression in these cases.

### Are Cajal bodies a checkpoint place before exporting previously retained mRNA into the cytoplasm?

In larch microsporocytes, a significant role in the process of nuclear retention and subsequent release of mRNAs into the cytoplasm should be attributed to Cajal bodies. From the very beginning of the cycle of cellular synthesis and turnover of poly(A) RNA, mRNAs are accumulated in them at 70-200x greater concentration than in the surrounding nucleoplasm. The highest level of mRNAs in CBs was observed just before the mass export of mRNAs to the cytoplasm, which indicates their relationship with switching the “retention program” to export (Figures 2A and 6A). This program change is probably related to the activation of mechanisms leading to the triggering of post-transcriptional splicing. In contrast to nucleoplasm, mature transcripts were not found in CB. Only non-fully-spliced transcripts with retained introns were stored in the Cajal bodies. This indicates that CBs are the place where a kind of switch must operate that releases retained transcripts from them and activates their post-transcriptional splicing.

To date, nuclear speckles (NSs) have been considered to be nuclear domains associated with pre-mRNA storage. It was shown recently that unspliced mRNAs (after SSA treatment), mRNAs with retained introns as well as completely spliced mRNAs were temporally sequestered in NS (Girard et al. 2012, Carvalho et al. 2017). In larch microsporocytes, the accumulation of polyadenylated transcripts in nuclear speckles appears to be at an incomparably lower level than in CBs (Kołowerzo et al. 2009). It remains unclear, however, why non-fully-spliced mRNAs are targeted to CBs. As it has already been shown, one of the reasons for the retention of transcripts containing intron sequences is the presence of spliceosome elements, such as U1 snRNA or U2AF, embedded in these sequences (Takemura et al. 2011). The reason for the retention of pre-mRNAs and associated spliceosomes in speckles is believed to be the presumable presence of speckle localization signal in all splicing factors (Girard et al. 2012). Thus, spliceosomes, composed of a multitude of splicing factors, might have a high affinity for speckles (Spector and Lamond 2011). Presumably, the strong affinity of splicing factors associated with non-fully-spliced mRNAs in microsporocytes determines their targeting to CBs. CBs function is most commonly associated with the production of spliceosomal enzymatic backbone, known as small nuclear RNPs, which catalyse RNA splicing. Both in early and late stage of snRNPs biogenesis, CBs are enriched in their components, precursors extended at their 3’ ends and fully assembled mature snRNPs (Frey et al. 2001, Stanêk and Neugebauer 2011, Wang et al. 2016). Additionally, CBs perform quality control on snRNPs through the U4/U6 di-snRNP recycling factor SART3 (Novotny et al. 2015). That’s why CBs may serve as “boosters” to aid the cell in spliceosomes assembly to accommodate to the additional load required in specific cells. Some research shows a correlation between the availability of spliceosome elements and aberrant splicing. The reduction in the production of functional snRNPs complexes may lead to widespread alternative splicing, especially IR events (Zhang et al. 2008).

In larch microsporocytes Cajal bodies appear to be an important element influencing the availability of splicing factors and the associated nuclear retention of mRNAs. On the one hand, they are a retention site for not fully spliced mRNAs encoding Sm proteins - a component of snRNP complexes. They also appear to be responsible for “turning on” the post-transcriptional splicing of these mRNAs, resulting in the formation of functional Sm proteins. On the other hand, they participate in the synthesis and assembly of snRNP elements and assembly of spliceosomal subunits. All this indicates that CBs can regulate the retention process of mRNAs other than Sm-encoding by regulating the rate of processing of individual elements of the snRNA complexes.

## STAR*methods

### Plant material and isolation of meiotic protoplasts

Larch (*Larix decidua* Mill.) anthers were collected from the same tree (Toruń, Poland, coordinates 53.021155 N, 18.570384 E) in successive meiotic prophase stages of diplotene (once a week from December 20, 2018 until January 20, 2019), to ensure constant experimental conditions. Anthers were fixed in 4% paraformaldehyde in phosphate-buffered saline (PBS), pH=7.2, for 12 h and then squashed to obtain free microsporocytes. Meiotic protoplasts were isolated from these cells according to Kołowerzo et al. (2009) and were then subjected to immunofluorescence-FISH reactions.

### Cytoplasmic fraction isolation and RNA purification

In order to obtain the cytoplasmic fraction for sequencing libraries generation, the plant material was homogenized in liquid nitrogen, and then the ribonucleoprotein extraction buffer was added in the proportion of 10 ml per 1.5 g of material (RNP buffer; 200 mM Tris-HCl pH 9.0, 110 mM potassium acetate, 0.5% (v / v) Triton X-100, 0.1% (v / v) Tween 20, 2.5 mM DTT, Roche protease inhibitor). After thawing on ice (~5 min), the material was filtered through a Miracloth monolayer (ø 22-25 μm; Merck Millipore) and centrifuged in a swing out centrifuge at 1500 × g for 2 min at 4°C. The obtained supernatant was the cytoplasmic fraction of the anther cells used for subjected to RNA purification and cDNA library preparation.

### Total RNA isolation

Total RNA isolation was performed either from cytoplasmic fraction (for sequencing libraries generation) or from suspension of microsporocytes (for PCR; freshly collected anthers were squashed into PBS buffer, and then the suspension was filtered on 70 μm sterile cell sieves, (BIOLOGIX Group) in order to obtain a pure fraction of microsporocytes; the procedure was performed on ice) using the Direct-zol ™ RNA Microprep (Zymo Research) according to manufacturer’s instructions.

### cDNA library preparation and sequencing

cDNA libraries generation and sequencing was performed using TruSeq Stranded Total RNA with Ribo-Zero Plant kit (Illumina) according to the protocol described in Hyjek-Składanowska et al. (2020).

### Reverse Transcription Reaction

RT was performed using SuperScript IV Reverse Transcriptase (Invitrogen) according to manufacturer’s instructions.

### PCR

The C1000 Touch Thermal Cycler (Bio-Rad) was used for all PCR reactions. The amplifications were performed in a final volume of 20 μl containing the following: 10 ng cDNA or 200 - 250 ng of genomic DNA, 1.5 mM MgCl_2_ (Syngen), 0.2 mM of each dNTP (Invitrogen), standard Gold Taq Buffer with 1 unit of Gold Taq DNA polymerase (Syngen) and 0.25 pM of each primer (sequences in Supl.1). The PCR cycling conditions for both primer pairs were as follows: denaturing at 95°C for 3 min, 34 cycles of 30 s at 95°C, 30 s at 56°C, 30 s at 72°C; final extension at 72°C for 5 min. The amplification products were analyzed using the HS DNA kit on Agilent 2100 BioAnalyzer according to manufacturer’s instructions.

### FISH in multiplex reactions

For hybridization, the probe was resuspended in hybridization buffer (30% v/v formamide, 4× SSC, 5× Denhardt’s buffer, 1 mM EDTA, 50 mM phosphate buffer) at a concentration of 50 pmol/ml. Hybridization was performed overnight at 26°C. The antisense DNA oligonucleotides (Genomed, Warsaw, Poland - sequences of the used oligonucleotides are included in Supplement 2) placed into a hybridization buffer at a concentration of 10 pmol/ml for each probe. Digoxygenine (DIG) probes were detected after hybridization using mouse anti-DIG (Roche) in 0.05% acetylated BSA in PBS (diluted 1:200) in a humidified chamber overnight at 11°C and anti-mouse Alexa 488 (Invitrogen) antibodies in 0,01% acetylated BSA in PBS (diluted 1:1000) in a humidified chamber for 1h at 36°C.

### Immunodetection of Sm proteins

For immunodetection, we used 0,1% NaBH_4_ for 2×5 minutes and wash with PBS, pH 7,2. Nonspecific antigens were blocked with PBS buffer containing 2% acetylated BSA for 15 minutes. Sm protein were detected by incubating with primary α-Sm antibody (ANA Human reference serum, Centers for Disease Control and Prevention, Atlanta GA 30333) in 0.1% acetylated BSA in PBS (1:300) in a humidified chamber overnight at 11°C and secondary anti-human antibody conjugated with Cy3, in 0.01% acetylated BSA in PBS (1:100) or and secondary anti-human antibody conjugated with Alexa 633 in 0.01% acetylated BSA in PBS (1:200) in a humidified chamber at 36°C for 1h.

### Design of multi-labeling reactions

Double-labeling immunofluorescence-FISH reactions (Sm proteins + mRNAs coding Sm proteins; Sm proteins + pol RNA II phospho CTD Ser2 + scaRNA) were performed as described above. In the double-labeling immunofluorescence-FISH reactions, the immunocytochemical methods always preceded the *in situ* hybridization methods because when *in situ* hybridization was applied first, the subsequent levels of immunofluorescence signals were very weak.

### Control reactions

For the immunofluorescence methods, control incubations lacking the primary antibody were performed. For the *in situ* hybridizations ribonuclease-treated samples were used. All control reactions produced negative results.

### Microscopy and quantitative measurements

The results were registered with Leica SP8 confocal microscope with an optimized pinhole, long exposure time (200 kHZ) and 63X magnification (numerical aperture, 1.4) Plan Apochromat DIC H oil immersion lens were used. Images were collected sequentially in blue (Hoechst 33342), green (Alexa 488), red (Cy3) and far red (Alexa 633, Cy5) channels. To minimize bleed-through between fluorescence channels, we employed low laser power (0.3% of maximum power) and sequential collection. For bleed-through analysis and control experiments, Leica SP8 software was used. Image acquisition was performed using constant parameters (laser power, detector gain, emission band and resolution). At least 6 cells from a given stage were analyzed. Three-dimensional optical sections were acquired with a 0.5 μm step interval. For all antigens and developmental stages, the obtained data were corrected for background autofluorescence, as determined by negative control signal intensities. ImageJ (Schneider et al. 2012) was used for image processing and analysis. Each reaction step was performed using consistent temperatures, incubation times and concentrations of probes and antibodies.

### Statistics

All statistical computations were performed in R 4.0.3 and significance level of 0.05 was used. Shapiro-Wilk’s and Levene’s tests (of the lawstat package) were used to check if normality and homogeneity of variance assumptions hold. As the assumptions did not hold, Kruskal–Wallis with Dunn’s post-hoc procedure (implemented in R package dunn.test) was applied in order to test for differences between multiple groups, i.e., signal levels at different stages. Benjamini-Hochberg false discovery rate correction was used. Spearman’s rank correlation test (rcorr function of the Hmisc package) was used to check if Sm proteins and their mRNA level in cytoplasm were correlated.

## Acknowledgements

This work was supported by Polish National Science Center NCN grant no. 2016/21/D/NZ3/00369 and grant NCN no 2015/19/N/NZ3/02410.

## Author contributions

M.R performed triple labeling of the RNA and proteins, PCR, confocal microscopy and image analysis, K.M, P.W.A performed triple labeling of the RNA and proteins, confocal microscopy and image analysis, M.S.H participated in experimental design, performed cDNA library preparation and participated in writing the MS, M.G. performed bioinformatical analysis and participated in discussion, analysis of results and MS writing, M.S performed confocal microscopy and image analysis and participated in discussion, A.J. designed experiments, participated in discussion and analyses of the results, D.J.S and A.K.L. designed experiments, performed triple labeling of the RNA and proteins, confocal microscopy and image analysis, participated in discussion and analyses of the results and writing the MS.

## Data availability statement

Data supporting the findings of this work are provided in the main text and the supporting information files. Raw reads generated during this project will be available at request from the corresponding authors. The full dataset for the transcriptomic analysis is available at https://repozytorium.umk.pl/handle/item/42. All other data and materials used in this study will be available at request from the corresponding authors.

